# PieParty: Visualizing cells from scRNA-seq data as pie charts

**DOI:** 10.1101/2020.08.25.267021

**Authors:** Stefan Kurtenbach, James J. Dollar, Anthony M. Cruz, Michael A. Durante, J. William Harbour

## Abstract

Single cell RNA sequencing (scRNA-seq) has been a transformative technology in many research fields. Dimensional reduction techniques such as UMAP and tSNE are used to visualize scRNA-seq data in two or three dimensions in order for cells to be clustered in biologically meaningful ways. Subsequently, gene expression is frequently mapped onto these plots to show the distribution of gene expression across the plots, for instance to distinguish cell types. However, plotting each cell with only one color leads to repetitive and unintuitive representations. Here, we present Pie Party, which allows scRNA-seq data to be plotted such that every cell is represented as a pie chart, and every slice in the pie charts corresponds to the gene expression of individual genes. This allows for the simultaneous visualization of the expression of multiple genes and gene networks. The resulting figures are information dense, space efficient and highly intuitive. PieParty is publicly available on GitHub at https://github.com/harbourlab/PieParty.

## Introduction

Gene expression data at single-cell resolution has brought a lot of opportunity to gain detailed understanding of heterogenous cell populations, but also many challenges regarding its processing and visualization. Principle component analysis (PCA) was initially used to reduce dimensions on single-cell RNA sequencing (scRNA-seq) datasets and to plot them in two or three dimensions. t-SNE and UMAP were developed subsequently, offering superior and global resolution, and are currently the most commonly used dimensional reduction techniques for scRNA-seq^1,2^. For most applications, gene expression is mapped onto t-SNE or UMAP plots, for instance to highlight different cell types or cell states. The common and only approach nowadays is to color the respective cell dots in UMAP or tSNE plots, where the intensity of the color correlates with how high the gene is expressed. However, this only allows to plot one gene per cell, which is inefficient, and results in many repetitive illustrations when expression of multiple different marker genes needs to be shown. This not only consumes valuable figure space, but is also not intuitive in many cases. Here we present PieParty, a python script that allows users to display each cell in t-SNE, UMAP, or any other single-cell plots with coordinates, as pie charts. Each pie chart can be used to visualize expression of multiple genes at once, and can be customized using different colors or color palettes. PieParty also offers additional settings for normalization and customized plotting.

## Results

The basic principle of the PieParty visualization is to generate pie charts for every cell in a single cell sequencing plot, like t-SNE and UMAP plots. The user provides a gene list of all genes that they want to visualize, and PieParty will generate pie charts where each slice in a pie chart represents the normalized gene expression of one individual gene in the list. Each gene (slice) can be assigned a unique color, or a color palette can be chosen to auto-assign unique colors for every gene. Choosing a color palette is useful for larger gene lists. As an example, we analyzed scRNA-seq data from human testis, and clustered the cells with UMAP^3^. The analysis reveals a developmental trajectory, ranging from stem cells to differentiated sperm, in very defined cell stages. Figure 1 and Supplementary Figure 1 shows PieParty plots, where every cell (n = 15,479) is represented by a pie chart, with 145 different differentiation markers plotted per pie chart. Each differentiation marker is automatically assigned a unique color on a color map, sorted from early markers in dark violet to markers of differentiation in yellow. The resulting plot shows that even plotting a large number of genes in pie charts still yields a very informative and intuitive UMAP plot, showing that the expression of these differentiation markers forms a continuum along the differentiation axis from stem cells to differentiated sperm cells.

**Figure 1:**
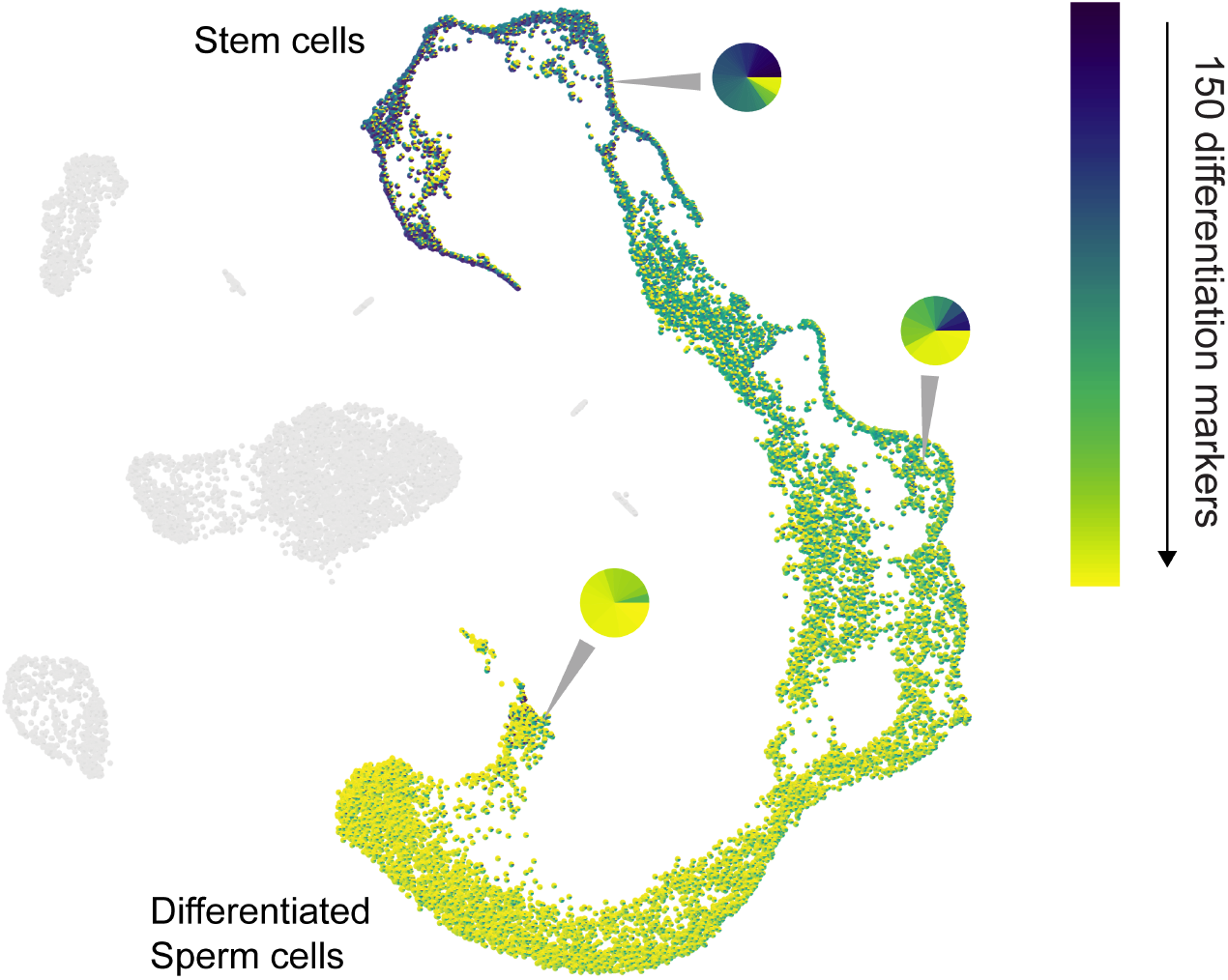
PieParty plot depicting scRNA-seq data from 5,479 testis cells. Around 150 differentiation markers were sorted from early (blue) to late (yellow), and auto-assigned with a color.

Besides merely plotting a single list of genes, PieParty also allows for more complex visualizations. As an example, we used scRNA-seq data from uveal melanoma (UM) tumors to demonstrate a more complex application^4^. There are two main classes of UM tumors, class 1, which rarely metastasize, and class 2, which frequently metastasize and have a high mortality rate. Eight biomarkers (EIF1B, FXR1, ID2, LMCD1, LTA4H, MTUS1, ROBO1, SATB1) are expressed in class 1 tumors, and four biomarker genes are up-regulated in the high-risk class 2 tumors (CDH1, ECM1, HTR2B, RRAB31). The classical way of visualizing this is by showing 12 individual UMAP plots. Supplementary Figure 2 shows all 12 genes plotted separately, which is very space consuming, and while still possible for 12 genes, not immediately comprehensible. For these and other cases, it is further useful to plot class 1 and class 2 tumor cells separately, which is depicted in Supplemental Figures 3 and 4. However, with 24 total plots this consumes even more precious space within a figure using the traditional way. PieParty offers several options to visualize this complex relationship of class 1 and class 2 genes. One consideration to make is that plotting two or more gene lists with different number of genes, in this case eight genes for class 1 versus four genes for class 2, comes with a bias which needs to be corrected for. If one gene list is twice the size of a second, but every gene is expressed in the same amount, the pie would be colored more with colors of the longer gene list. Hence, PieParty allows to normalize for that fact, and weighs gene sets equally by applying a normalization factor to account for the difference in number of genes in each gene list. This normalization technique was applied for all following plots, and the color intensity of the individual pie slices was set to correlate with gene expression, in addition to the size of the pies itself.

Figure 2a depicts the tSNE plots of 11 UM tumor samples, (3 class 1, 8 class 2, n = 59,915 cells) plotted with PieParty and a simple blue/red color scheme. Class 1 and Class 2 tumor cells were plotted individually, as well as combined in one big plot. It is immediately evident, that the tumor cell markers are not only found in tumor cells, but also in immune cells. Class 2 tumor markers are predominantly enriched in macrophages and monocytes, whereas class 1 tumor markers are present in other immune cells including Lymphocytes, NK cells, T-Cells. Especially interesting is the fact that in this scRNA-seq dataset one class 1 tumor marker is expressed in some class 2 immune cells. While all this information is technically visible when all genes are plotted individually in 24 plots (see Supplemental Figures 3 and 4), the PieParty plots are intuitively readable and visualize these relationships in a space efficient way. The information density can be further increased by assigning individual colors to each gene, which PieParty can do automatically. Figure 2b shows the same dataset with color palettes applied automatically to the two gene lists. All labels indicating which gene was assigned which colors are generated by PieParty. This visualization shows that for class 2 tumors the biomarkers expressed by immune cells are different from the genes expressed by tumor cells, whereas for class 1 tumors there is overlap. Together, visualizing this dataset with PieParty, proportional expression of biomarkers, as well as expression amount for each gene are all assessable in one single plot.

**Figure 2:**
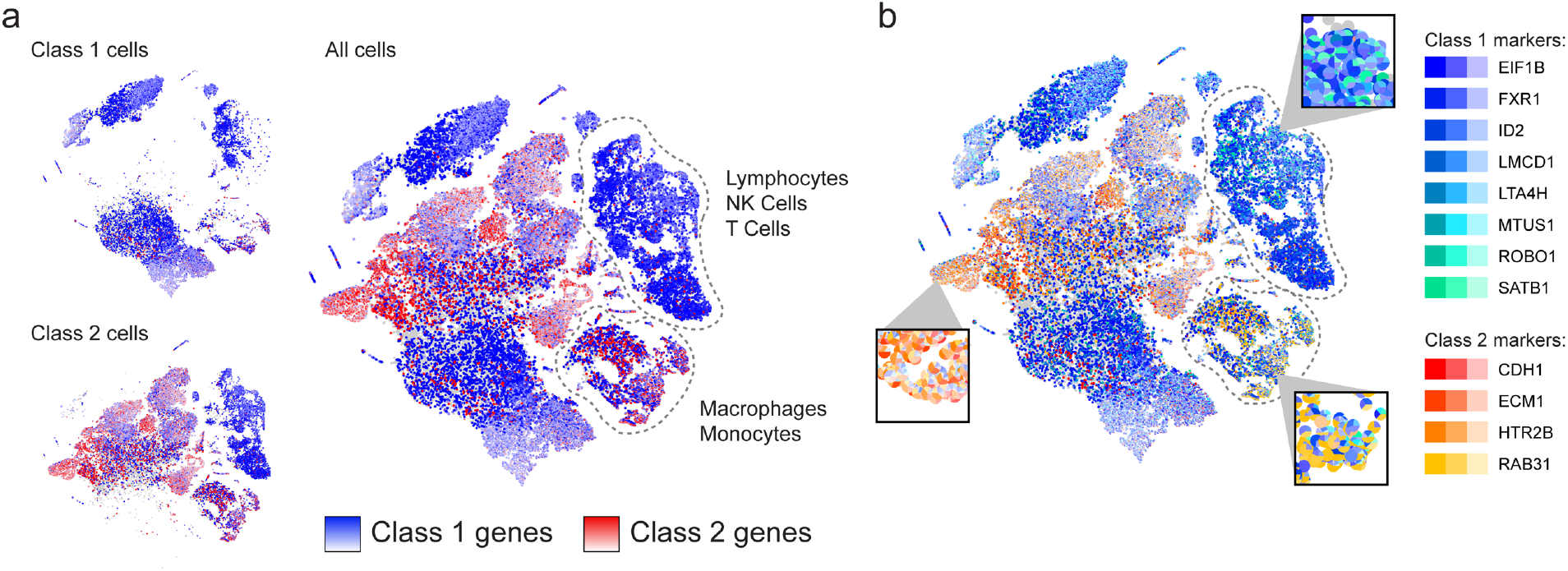
(a) Class 1 and Class 2 tumor cells plotted separately, with class 1 associated genes in blue, and class 2 associated genes in red. The big plot on the right side of (a) combines all class 1 and class 2 cells (“All cells”) (b) Same plot as in Figure 2a, but with auto-assigned color palettes, giving each gene a unique color. Color intensity of each slice correlates with gene expression, in addition to slice size.

Another field of application for PieParty is distinguishing cell types and identifying rare cell populations. As an example, we visualized different immune cells from the UM dataset. Figure 3a shows the macrophage population extracted from the complete dataset (n = 8,048 cells), where three M1 macrophage markers assigned the color red, and eleven M2 macrophage markers were assigned blue. As mentioned above, PieParty normalized by gene numbers in the different marker lists, which permits for lists with very different number of genes to be utilized like in this case. Figure 3a shows the clear visualization of M1 and M2 macrophages on the right side of the plot, strongly expressing the indicated markers. Interestingly, there is an equally big population of cells present that express mainly M2 markers but weakly, which cluster to the left of the M1 and M2 cells. This provides a striking and clear visualization of macrophage activation heterogeneity and phenotype diversity within the tumor immune microenvironment, Hence, this visualization allows to distinguish these very similar cell types in one plot in a data-driven way, and display the landscape of the expression data. This approach can also be applied to highlight rare cell populations. As an example, we extracted macrophages, monocytes, and undetermined lymphocytes form the UM scRNA-seq dataset (n = 5,111 cells) (Figure 3b). In this dataset T-cells are a rare cell population, comprising only 1.1% of the total cells. We distinguished T-cells by expression of CD3G, CD4, CD8A, and CD8B from monocytes and macrophages, which express CD14, FCGR3A, FCGR1A, CD68, TFRC, CCR5, and ITGAM. The PieParty plot shows how a rare cell population can be clearly distinguished directly in a data-driven way, in contrast to relying on a manually labeled cell cluster.

**Figure 3:**
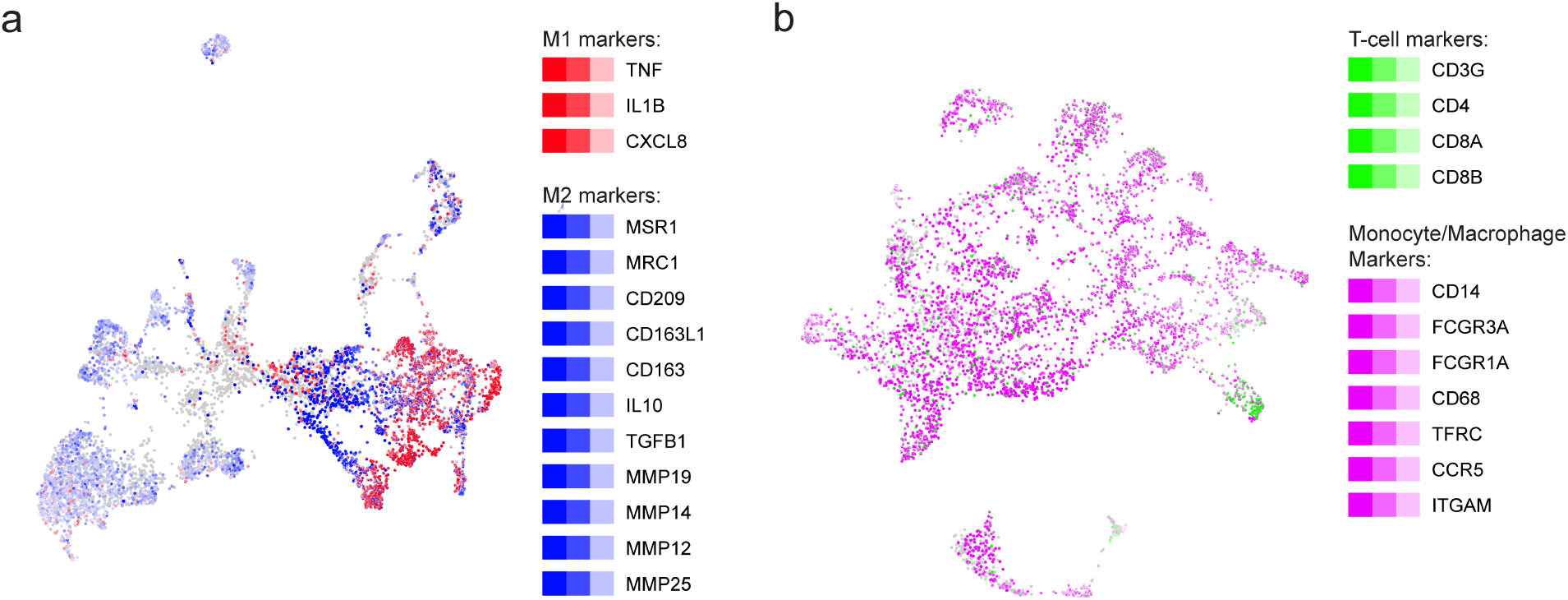
PieParty plots of immune cell populations of uveal melanoma tumors. (a) M1 (red) and M2 (blue) macrophage markers are plotted to distinguish M1 and M2 macrophages. (b) Use pf PieParty to plot rare cell populations distinguished by multiple markers. T-cell markers (green) are combined to highlight this rare cell population and distinguish them from monocytes and macrophages. Color intensities correlate with gene expression levels.

Another functionality in PieParty is to plot the average gene expression per cell cluster. Figure 4 depicts the “plot clusters” functionality, which generates one pie chart per cell cluster, with the pie chart size correlating with the number of cells in the respective cluster. This plotting style can greatly simplify complex datasets, and allow for an intuitive assessment of the expression of different markers in the clusters.

**Figure 4:**
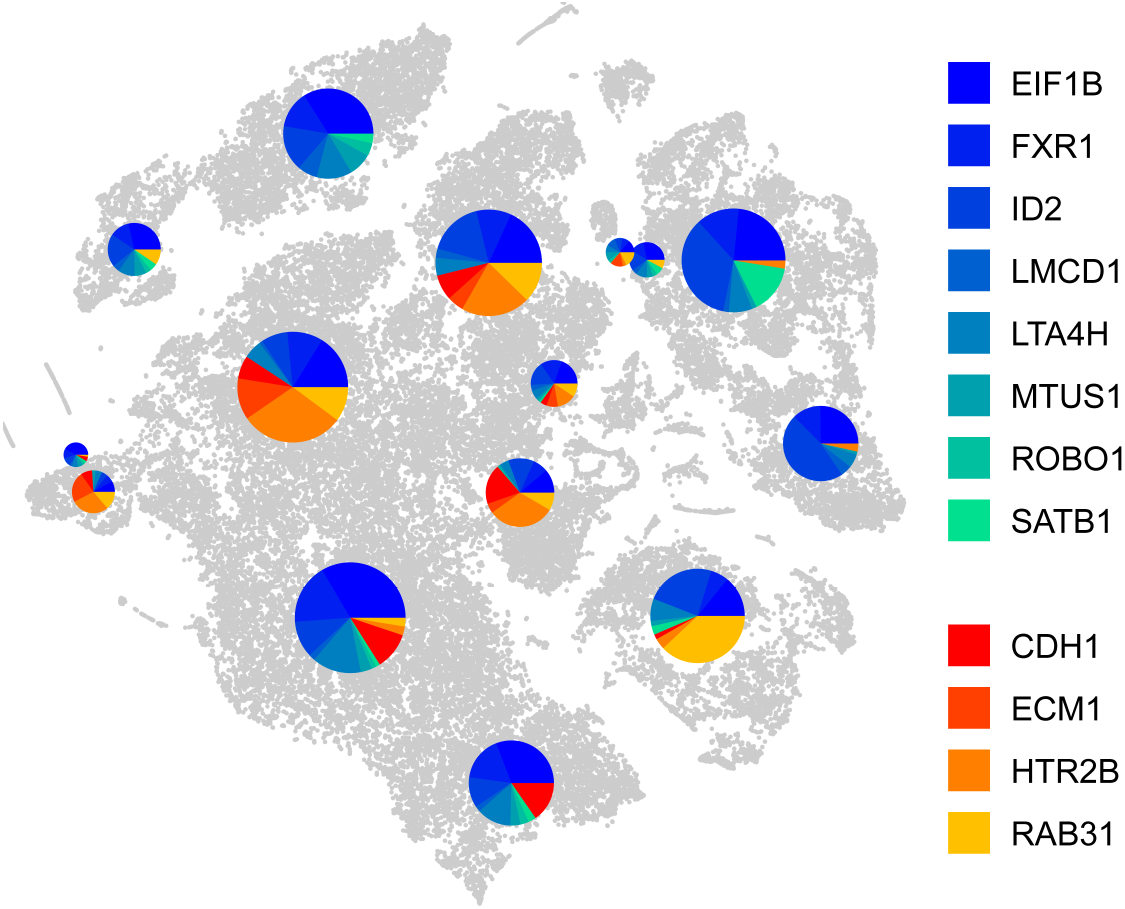
PieParty cluster plot. Each cell cluster is assigned one pie chart depicting the average gene expression in the respective cluster. The chart sizes correlate with the number of cells in the cluster. Class 1 tumor genes were auto-assigned colors from the colormap “winter” (blue-green), and class 2 tumor genes were colored with “autumn” (red-yellow).

## Discussion

Here we present PieParty, which allows single cells from scRNA-seq data to be represented as pie charts instead of single-colored dots. PieParty plots are far more information rich, allowing multi-dimensional single cell sequencing data to be represented in an intuitive and space-efficient way, and for the identification of previously unrecognized transcriptional heterogeneity. We have presented use cases including gene lists with large number of genes to display expression of differentiation markers in sperm differentiation (Figure 1), discern tumor risk classes (Figure 2), characterize immune cell infiltrates (Figure 3a), as well as highlight rare cell populations (Figure 3b). With the public availability of various gene sets, this new visualization technique can be utilized to visualize various cell types, cell states, pathways, cell cycle, senescence, gene networks and many more^5^. Together, PieParty can provide deeper insights into biology derived from single cell sequencing data by allowing for multidimensional visualization of high-density datasets.

## Methods

scRNA-seq data for testis and UM are publicly available ^3,4^, and were downloaded, analyzed with Seurat (Version 3.2.2) as previously described^4^. The pure macrophage population was generated by extracting cells with CD68 expression > 1. The macrophages, monocytes, and undetermined lymphocytes population used for rare cell population identification was generated by extracting cell clusters based on cell type classifications generated from the original analysis^4^.

PieParty plots were generated using PieParty 1.4 - 1.8, with standard settings. For Figure 1a “lighten colors”, “-lc” was set to “False”.

## Author contribution statement

SK developed PieParty, and prepared the manuscript. JJD helped with design, data acquisition, analysis, and manuscript preparation. MAD helped with design and manuscript preparation. JWH helped with overall design, data interpretation, and manuscript preparation.

## Acknowledgements

This work was supported by Melanoma Research Foundation Career Development Award (Kurtenbach) and Established Investigator Award (Harbour), National Cancer Institute grant R01 CA125970 (Harbour), A Cure in Sight Jack Odell-John Dagres Research Award (Kurtenbach, Harbour), Bankhead-Coley Research Program of the State of Florida (Harbour), The Helman Family-Melanoma Research Alliance Team Science Award (Harbour)and a generous gift from Dr. Mark J. Daily (Harbour). The Bascom Palmer Eye Institute received funding from NIH Core Grant P30EY014801 and a Research to Prevent Blindness Unrestricted Grant. The Sylvester Comprehensive Cancer Center also received funding from the National Cancer Institute Core Support Grant P30CA240139. The content is solely the responsibility of the authors and does not necessarily represent the official views of the National Institutes of Health.

---

**Supplementary Figure 1:**
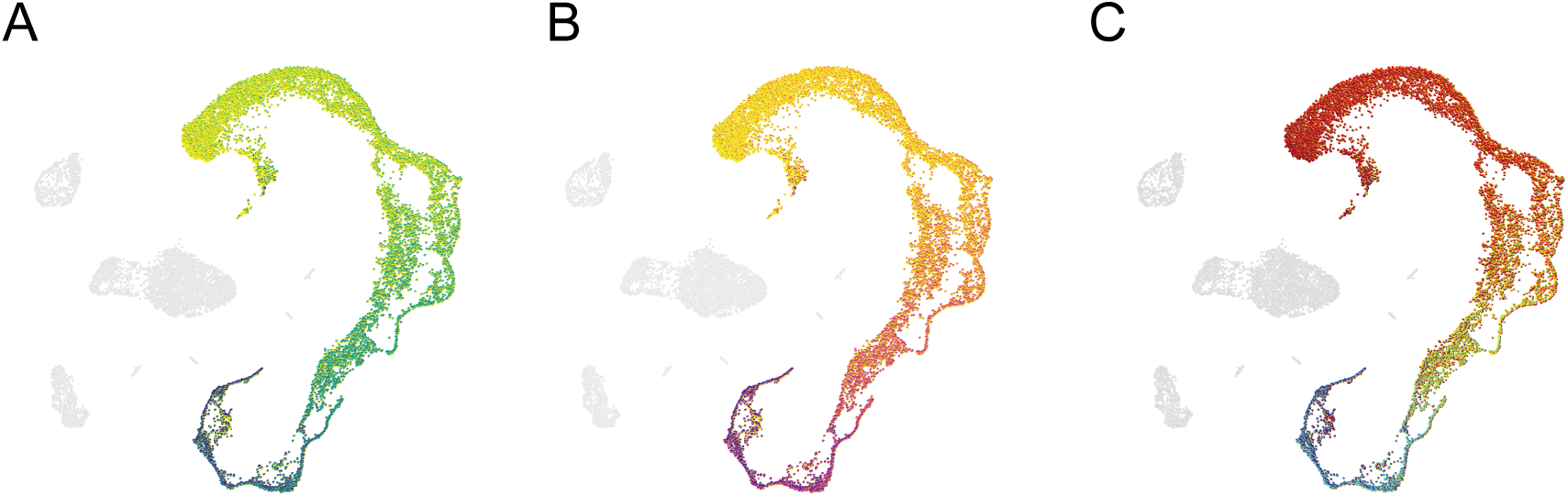
Different color maps applied to the data shown in Figure 1. The colors are automatically assigned to individual genes.

**Supplementary Figure 2:**
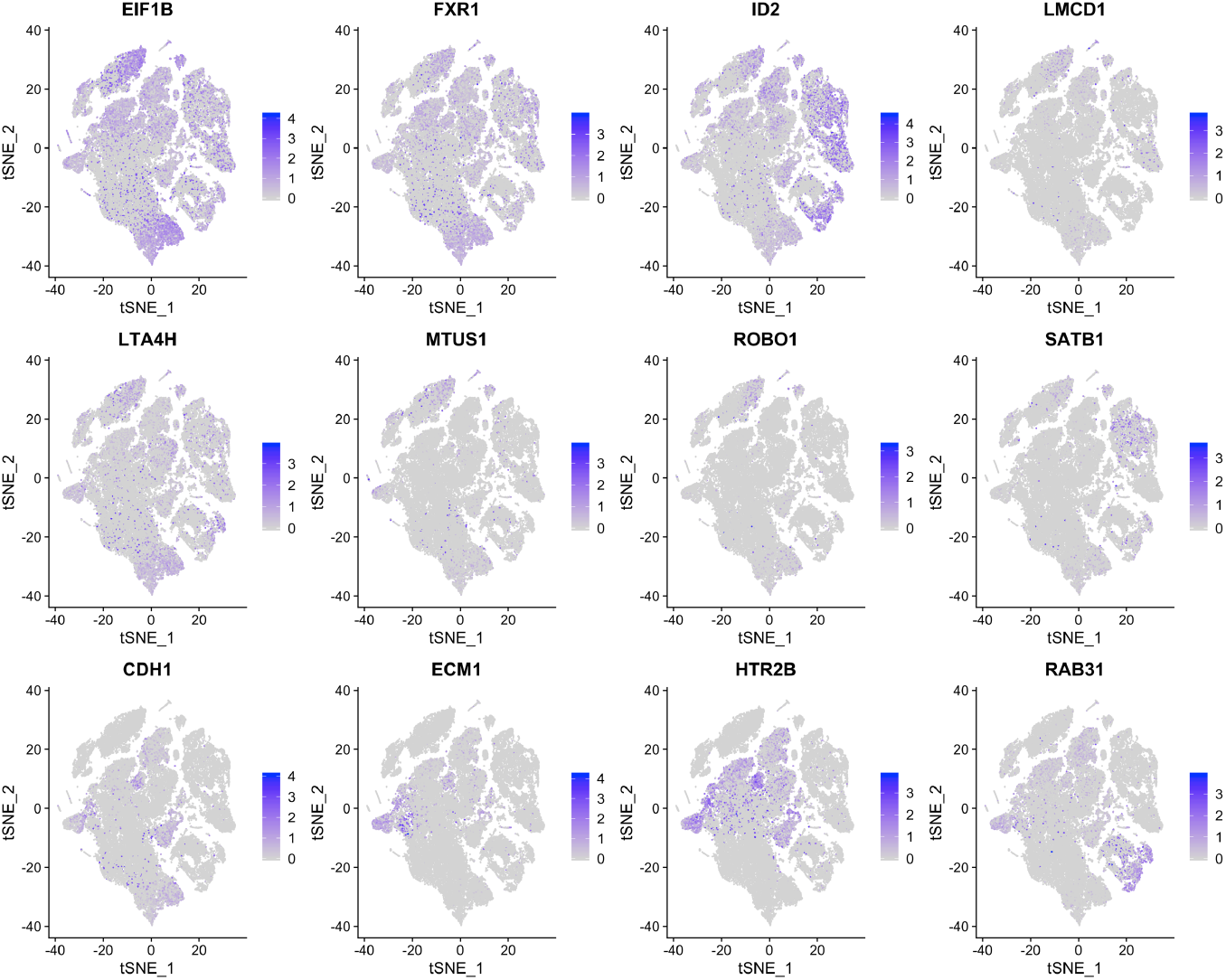
tSNE plots for twelve uveal melanoma biomarker genes.

**Supplementary Figure 3:**
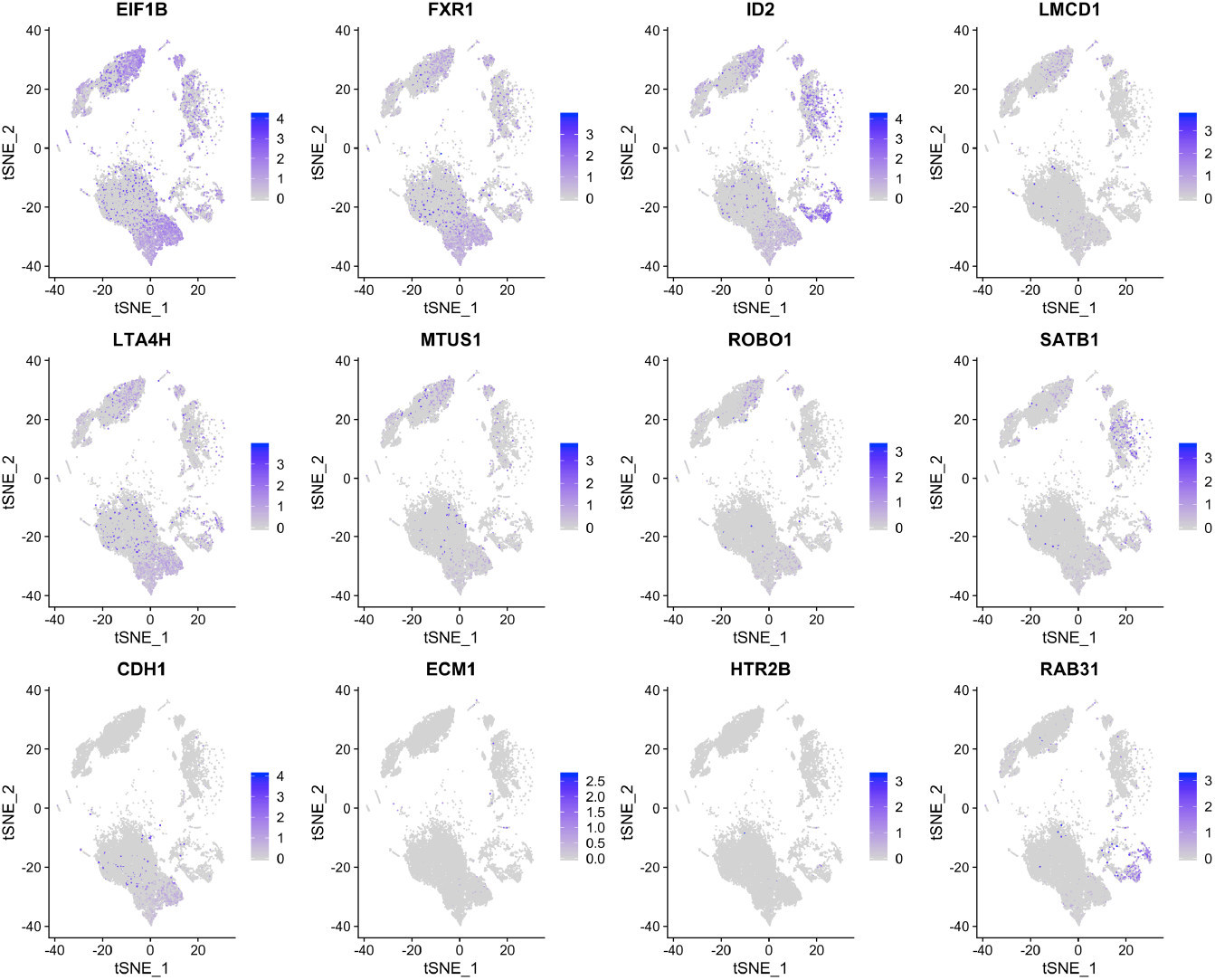
tSNE plots for twelve uveal melanoma biomarker genes in class 1 uveal melanoma cells extracted from the entire dataset.

**Supplementary Figure 4:**
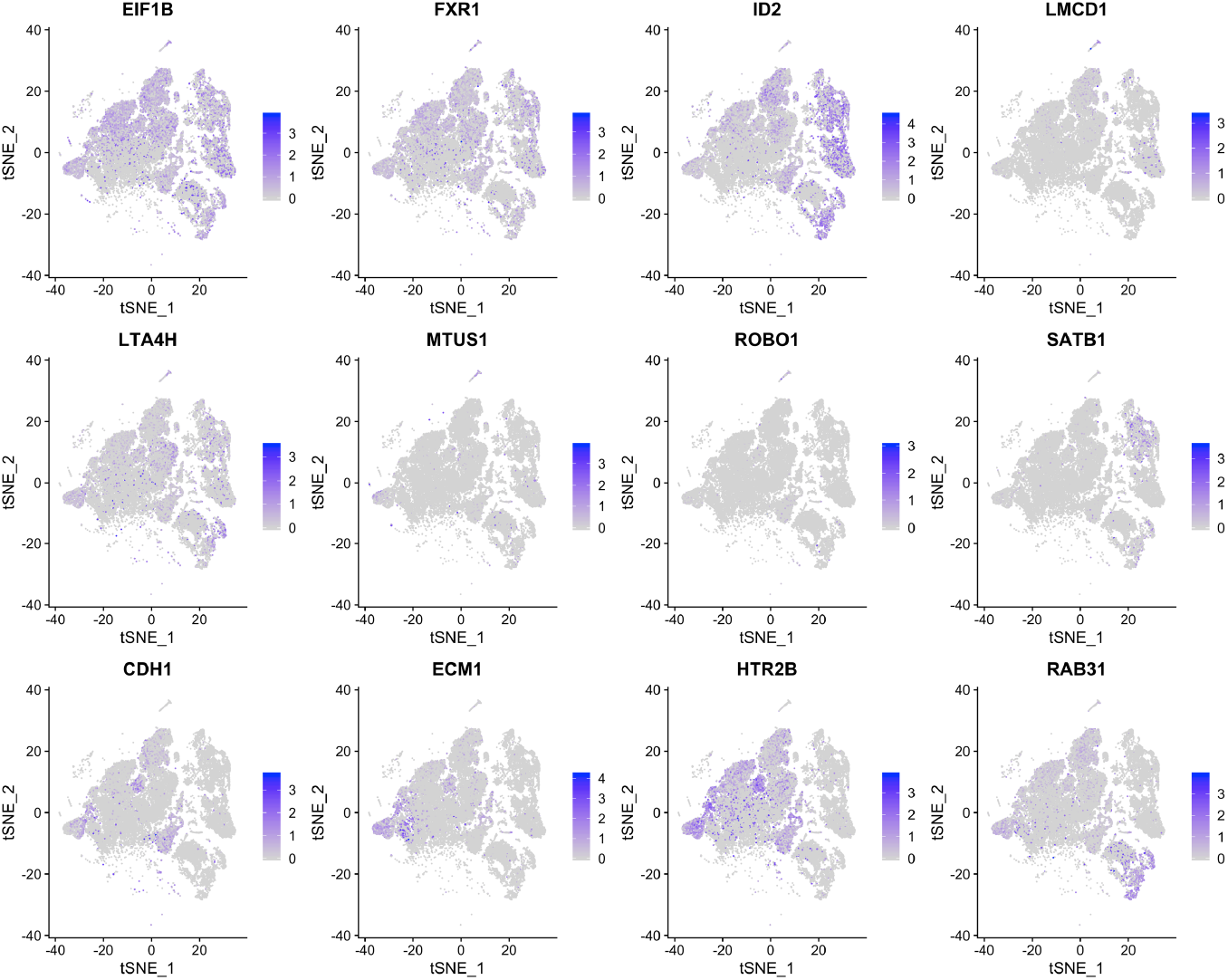
tSNE plots for twelve uveal melanoma biomarker genes in class 2 uveal melanoma cells extracted from the entire dataset.

## Notes

https://github.com/harbourlab/PieParty

### Competing Interest Statement

The authors have declared no competing interest.

## References

1. van der Maaten, L. & Hinton, G. Viualizing data using t-SNE. Journal of Machine Learning Research 9, 2579–2605 (2008).

2. McInnes, L., Healy, J. & Melville, J. Umap: Uniform manifold approximation and projection for dimension reduction. arXiv preprint arXiv:1802.03426 (2018).

3. Guo, J. et al. The adult human testis transcriptional cell atlas. Cell Res 28, 1141–1157, doi:10.1038/s41422-018-0099-2 (2018).

4. Durante, M. A. et al. Single-cell analysis reveals new evolutionary complexity in uveal melanoma. Nat Commun 11, 496, doi:10.1038/s41467-019-14256-1 (2020).

5. Subramanian, A. et al. Gene set enrichment analysis: a knowledge-based approach for interpreting genome-wide expression profiles. Proc Natl Acad Sci U S A 102, 15545–15550, doi:10.1073/pnas.0506580102 (2005).

